# Metabolic Energy Sensing by Mammalian CLC Anion/Proton Exchangers

**DOI:** 10.1101/545368

**Authors:** Matthias Grieschat, Katharina Langschwager, Raul E. Guzman, Christoph Fahlke, Alexi K. Alekov

**Author notes:** Corresponding author: Alexi K. Alekov, Ph.D., Institute of Neurophysiology, Hannover Medical School, OE4230, Carl-Neuberg-Str. 1, 30625 Hannover, Germany. These authors contributed equally to this work (Matthias Grieschat, Katharina Langschwager).

## Abstract

Mammalian CLC anion/proton exchangers control the pH and [Cl^-^] of the endolysosomal system, one of the major cellular nutrient uptake pathways. We explored the regulation of the vesicular transporters ClC-3, ClC-4, and ClC-5 by the adenylic system components ATP, ADP, and AMP. Using heterologous expression and whole-cell electrophysiology, we demonstrated that cytosolic ATP and ADP but not AMP and Mg^2+^-free ADP enhance CLC ion transport via binding to the protein C-terminal CBS domains. Biophysical investigations revealed that the effects depend on the delivery of intracellular protons into the CLC transport machinery and result from modified voltage-dependence and altered probability that CLC proteins undergo silent non-transporting cycles. Our findings demonstrate that the CLC CBS domains are able to serve as energy sensors by detecting changes in the cytosolic ATP/ADP/AMP equilibrium. The adenine nucleotide regulation of vesicular Cl^-^/H^+^ exchange creates a link between the activity of the endolysosomal system and the cellular metabolic state.

## Introduction

CLC anion channels and anion/proton exchangers fulfill indispensable functions in nerve and muscle excitation, endocytosis, exocytosis, and lysosomal function [1,2]. CLC proteins assemble as dimers of two identical subunits with parallel orientation and separate ion transport pathways. The intracellular carboxy terminus (C-terminus) of each subunit contains a so-called Bateman domain with two distinct cystathionine β-synthase domains, CBS1 and CBS2 [3–7]. CBS domains are adenine-nucleotide-binding structures found in many unrelated protein families; their physiological function has been established by association with various human hereditary diseases [8–11]. A good example are the proteins of the AMPK family (AMP-activated protein kinases) in which the CBS domains serve as energy sensors detecting changes in the ATP/ADP/AMP levels (see [12] for a review).

Mammalian CLC transporters (ClC-3 to ClC-7) are localized mainly on intracellular vesicles and regulate the vesicular acidity and Cl^-^ content [2]. Presently, only the structure of the ClC-5 C-terminus is described revealing an adenine nucleotide binding pocket formed between the two cytoplasmic CBS domains and occupied either by ADP or ATP ([6], Figure 1A). To our knowledge, the AMP-occupied and the nucleotide-free CBS domain conformations have not been characterized yet for any of the mammalian CLC transporters. Nevertheless, large-scale ATP-induced conformational changes of the ClC-5 C-terminus demonstrate unequivocally the regulatory capacity of the nucleotide binding pocket [13]. Combined, the above data suggest that mammalian CLC transporters are able to detect and respond to changes in the cellular energy status; however, the functional effects of nucleotide binding are insufficiently understood. Controversial findings have been reported on the very similar isoforms ClC-4 and ClC-5 [14,15]. Whereas ClC-4 was found to discriminate between ATP and ADP [14], ClC-5 was reported to behave similarly in the presence of ATP, ADP and AMP [15]. Of note, the release or production of metabolic energy results predominantly in ATP/ADP/AMP interconversion and a nearly constant total cellular content of these nucleotides [12]. Hence, the different effects of adenine nucleotides on ClC-4 suggest that the CLC CBS domains may serve as energy sensors [14]. In contrast, the identical functional effects of ATP, ADP and AMP on ClC-5 implicate that nucleotide binding in mammalian CLC transporters is physiologically irrelevant [15].

**Figure 1.**
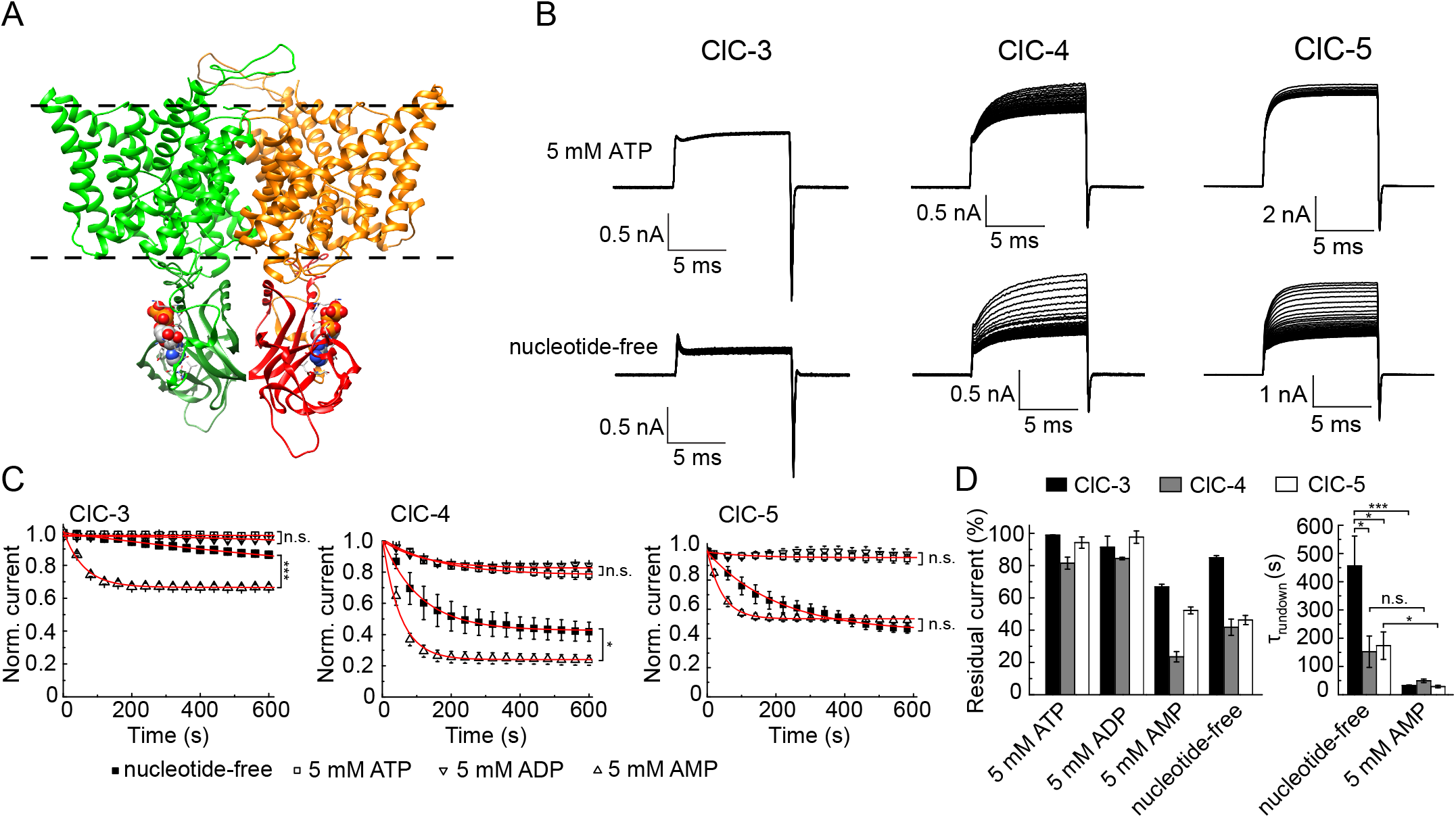
Cytosolic adenine nucleotides regulate the function of ClC-3, ClC-4, and ClC-5. **(A)** Homology model depicting the ClC-5 dimeric structure with bound ATP molecules represented as spheres. Some of the nucleotide-coordinating residues between in the C-terminus CLC CBS domains are represented as sticks. Dashed lines indicate the hydrocarbon boundaries of the lipid bilayer. **(B)** Representative leak-subtracted ClC-3, ClC-4 and ClC-5 whole-cell currents measured with (upper row) or without (lower row) 5 mM ATP added to the pipette solution. Currents were elicited by voltage steps from 0 mV to +140 mV, followed by a step to −100 mV to maximize the off-gating currents visible as sharp peaks at the end of the pulses. The recording started shortly (10s - 30s) after obtaining a whole-cell configuration. NMDG-based solutions were used to characterize ClC-4. **(C)** Effects of various adenine nucleotides or their washout on the ionic transport of ClC-3, ClC-4 and ClC-5. The current amplitudes were measured at the end of the test pulses from experiments as shown in *(B)* and normalized to the initial current amplitudes obtained after establishing the whole-cell configuration. Some of the experimental data points have been omitted for clarity. Monoexponential fits to the data are included as red lines. Asterisks indicate statistical significances obtained by t-test analysis of the residual currents determined at the end of each 10-minute measurement (*, p < 0.05; ***, p < 0.001, n.s., not significant; n= 5-11). **(D), Left panel:** residual currents determined at the end of 10-minute measurements as shown *(C)*. **(D), Right panel**: time constants (τ) of the current rundown obtained by fitting monoexponential functions to individual measurements as depicted in (C). Results from the t-test analysis of the data are indicated as described in (C).

To clarify whether mammalian CLC transporters can sense and respond to metabolic energy changes, we reinvestigated the effects of ATP, ADP and AMP on ClC-4 and ClC-5. In addition, we tested whether adenine nucleotides can functionally regulate the neuronal transporter ClC-3. Our data revealed that all investigated CLC isoforms can discriminate between cytosolic ATP and AMP and are, therefore, able to detect changes in the cellular energy status. Biophysical analysis demonstrated that the molecular mechanisms of adenine nucleotide regulation are complex and target the voltage dependent activation of the mammalian CLCs, as well as their proton transport machinery.

## Results

### The ion transport of ClC-3, ClC-4, and ClC-5 is differentially regulated by intracellular ATP, ADP and AMP

We transfected HEK293T cells and used whole-cell electrophysiology to investigate the functional effects of adenine nucleotides on three CLC anion/proton exchanger isoforms, the human ClC-4 and ClC-5, and the membrane-localized mouse ClC-3 splice variant ClC-3c [16]. The expression resulted in large macroscopic currents with the specific biophysical properties of the corresponding isoforms (Figure 1B) [17]. In whole-cell mode, the content of the cytoplasm is exchanged for the pipette solution by diffusion [18]. Hence, tracking the CLC current amplitude after establishing whole-cell configuration can be used to detect effects resulting from washout of endogenously present cytosolic nucleotides or from their substitution by nucleotides added to the pipette solution. To mimic the physiological conditions, we always added 5 mM MgCl_2_ into the pipette solution (unless explicitly stated otherwise).

Experiments with 5 mM ATP in the pipette solution revealed a minor CLC transport decrease (Figure 1B-D). As previously demonstrated [14,15], a much larger CLC transport decrease was observed for all investigated isoforms in experiments with nucleotide-free pipette solution (Figure 1B-D). These results suggest that intracellular ATP binding enhances the ion transport of ClC-3, ClC-4 and ClC-5. To support this hypothesis, we tested the reversibility of the effects by applying ATP to the internal side of excised inside-out patches from cells expressing ClC-5. The application reversibly increased the CLC current amplitude (Supplementary Figure S1). Therefore, the current rundown induced by the washout of cytoplasmic ATP does not result from internalization of CLC proteins but from reduced CLC ion transport. Next, we mutated one of the amino acids coordinating the ATP molecule in the ClC-5 nucleotide binding pocket (mutation Y617A, [6]). As expected, the mutant abolished the current decrease associated with the washout of cytosolic ATP (Supplementary Figure S2). The result demonstrates the role of the C-terminal CBS domains in the adenine nucleotide regulation of mammalian CLC transport.

To test whether mammalian CLC transporters can detect metabolic energy changes, we performed analogous experiments with intracellular ADP and AMP. The ionic transport of the investigated isoforms was not substantially reduced when 5 mM ADP was added to the pipette solution. In contrast, a large reduction was observed with 5 mM AMP in the pipette. For ClC-3 and ClC-5, the AMP-induced rundown was significantly faster compared to nucleotide-free measurements (Figure 1C, 1D). In addition, the residual currents in experiments with AMP-containing pipette solution differed from nucleotide-free measurements (note the different steady-state amplitudes reached at the end of the 10-minute measurements). These two observations suggest that AMP can displace ATP from the C-terminal CLC binding pocket. To test this hypothesis, we conducted experiments with both 1 mM AMP and 5 mM ATP in the pipette solution. In agreement with the proposed competitive displacement of ATP by AMP (see also [13,20]), the ion transport decrease in these experiments was larger than the one in experiments conducted with only ATP present in the pipette (Supplementary Figure S3).

In summary, our findings demonstrate that cytosolic ATP and ADP increase the ion transport rates of ClC-3, ClC-4, and ClC-5. In contrast, we found that cytosolic AMP decreases CLC ion transport. The different magnitudes of the effects suggest, further, that adenine nucleotide regulation might be optimized to match the physiological roles of the specific CLC isoforms.

### Cytosolic adenine nucleotides regulate the CLC transport cycle and voltage-dependent gating

Cytosolic adenine nucleotides regulate the ion currents of the CLC channels ClC-1 and ClC-2 by altering their voltage-dependent activation [21–23]. Similar to CLC channels, the here-investigated CLC transporters also exhibit pronounced voltage dependence [24,19,17]. Therefore, we used gating current analysis [17,24,25] to test whether adenine nucleotides alter the voltage dependent activation of ClC-5. Specifically, we calculated the gating charge using the area beneath the gating currents of this transporter (Figure 2A). The analysis revealed that ATP, ADP and AMP all shift the ClC-5 activation towards more positive voltages (Figure 2B, Supplementary Figure S4A, Supplementary Table S1). This shift may contribute to the AMP-induced current decrease; however, it cannot explain the ClC-5 transport increase observed with ATP and ADP in the pipette solution.

**Figure 2.**
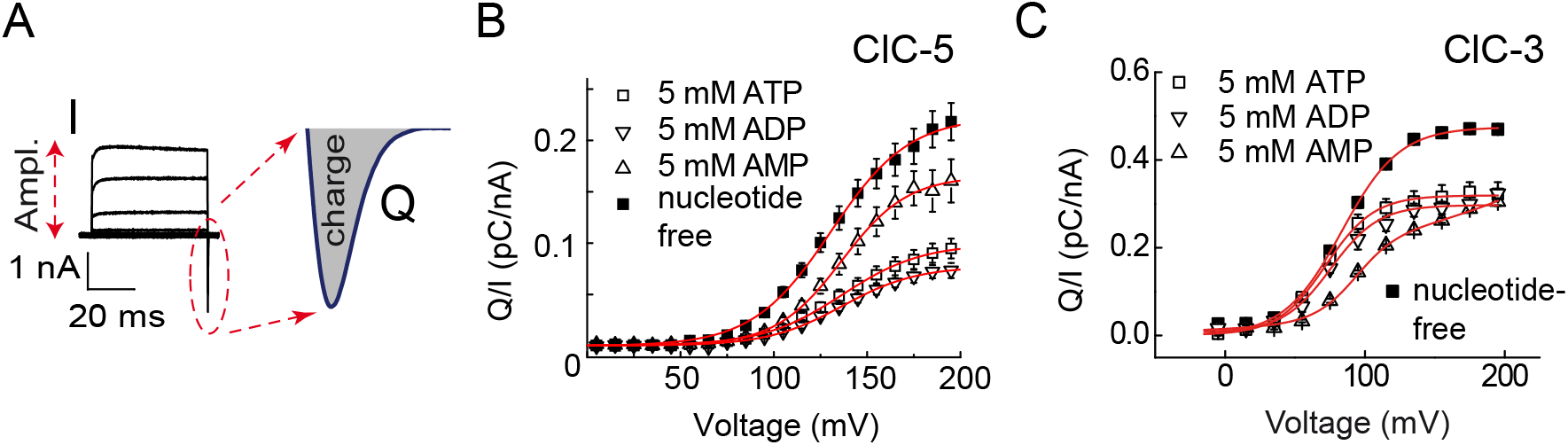
Adenine nucleotides regulate the CLC voltage dependence and silent transport cycle probability. **(A)** Schematic representation of the gating charge analysis. The gating charge Q was obtained by calculating the surface beneath the off-gating currents (enlarged in gray and denoted by “charge”). The apparent silent transport cycle probability (see (B)) was obtained by dividing the gating charge by the ion current amplitude I (“Ampl.”) at +165 mV. **(B)** Voltage dependence of the WT ClC-5 off-gating charge normalized to the ionic current at +165 mV in the absence or presence of adenine nucleotides. Red lines represent Boltzmann fits to the data; fit parameters are provided in Supplementary Table S1. **(C)** Voltage dependence of the WT ClC-3 off-gating charge normalized to the ionic current at +165 mV in the absence or presence of adenine nucleotides (n=4-5). Red lines represent Boltzmann fits to the data; fit parameters are provided in Supplementary Table S2.

It was previously established that impaired delivery of internal protons into the CLC transport machinery drives mammalian CLC exchangers towards a silent non-transporting mode with reduced ion transport but increased gating current amplitudes [17,24,25]. Hence, the ratio between the gating charge and the ion current at one fixed voltage can be used to estimate the number of silent non-transporting CLC proteins (see Figure 2A). Applying this analysis to ClC-5 revealed that ATP and ADP both decrease the aforementioned ratio (Figure 2A and 2B). Accordingly, these nucleotides seem to enhance CLC ion transport by reducing the probability that ClC-5 undergoes silent non-transporting cycles. In contrast, AMP binding increased the ratio and accordingly the silent transport cycle probability. Of note, the apparent ClC-5 gating charge in experiments with 5 mM ATP differed from the one measured with both 5 mM ATP and 1 mM AMP added to the pipette solution (Supplementary Figure S4B). This difference provides additional evidence that AMP can displace ATP from the CBS nucleotide-binding pocket.

Analyzing the gating currents of ClC-3 revealed that cytosolic adenine nucleotides also affect both the voltage dependence and the silent transport cycle probability (Figure 2C). However, in contrast to ClC-5, cytosolic AMP induced a stronger right-shift of the ClC-3 voltage-dependent activation. This explains well the different responses of ClC-3 and ClC-5 to AMP when compared to measurements with nucleotide-free pipette solution (Figure 1), and hints at the physiological specialization of these CLC transporters.

### The proton transport pathway is important for CLC adenine nucleotide regulation

Voltage dependent gating in CLC channels relies upon a conserved negatively charged glutamate residue at the extracellular entrance of the anion transport pathway, the so-called gating glutamate Glu_ext_ (see [4] for a review). Glu_ext_ plays also a critical role in the voltage-dependent gating of ClC-3, ClC-4 and ClC-5 [26–28,24,29]. In addition, Glu_ext_ constitutes a critical part of the CLC proton transport pathway [26,27,30]. This led us to evaluate the specific role of Glu_ext_ in the adenine nucleotide regulation of mammalian CLC transporters. To this end, we characterized the effect of cytosolic ATP on the pathogenic mutant E211G that neutralizes the electric charge of the Glu_ext_ E211 and abolishes the depolarization-induced activation of ClC-5 [31]. Experiments with nucleotide-free pipette solution revealed that ATP depletion reduces the ion transport of E211G ClC-5 (Figure 3A, 3B, Supplementary Figure S5). Thus, the Glu_ext_-dependent activation process is not the sole determinant of the CLC adenine nucleotide regulation.

**Figure 3.**
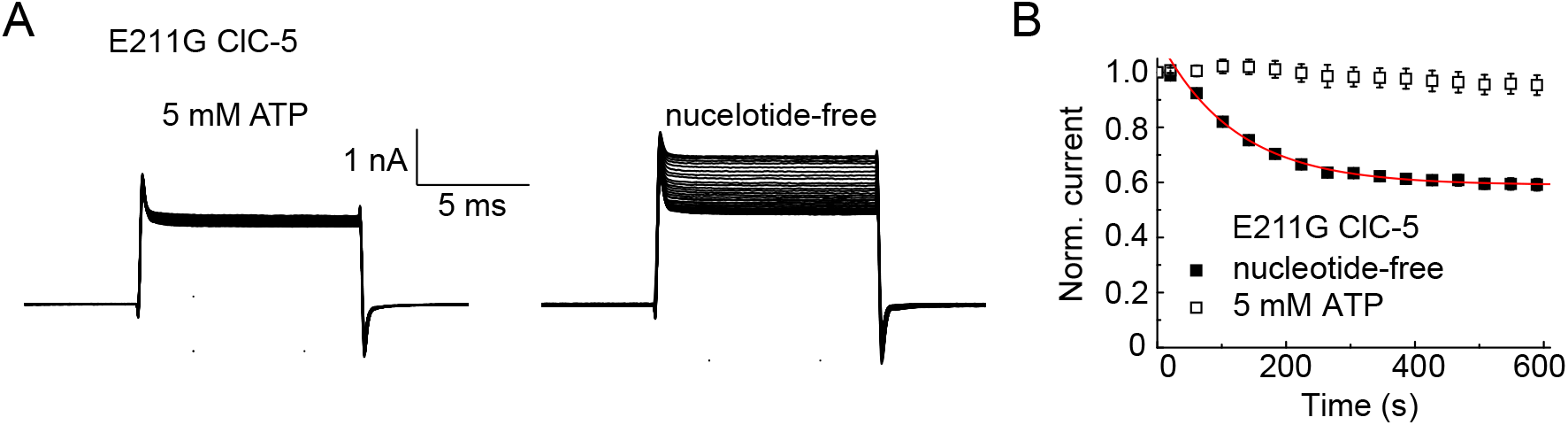
Effects of nucleotide chemistry and Mg^2+^. **(A)** Representative whole-cell recordings of cells expressing E211G ClC-5 upon voltage pulses from a −60-mV holding potential to +140 mV applied in 10-s intervals starting shortly after obtaining the whole-cell configuration with (left) or without (right) 5 mM ATP in the pipette solution. Asymmetric [Cl^-^] was used (110 mM internal NaCl substituted by TrisSO_4_) to achieve a stable negative reversal potential of <-50 mV allowing to exclude recordings with high unspecific leak current contamination. Leak subtraction was not applied. (**B**) Normalized time course of the steady-state E211G ClC-5 current amplitudes measured with (n=5) or without (n=4) internal ATP. Red line represents a monoexponential fit to the data with a time constant of 108±8 s; currents measured with ATP-free pipette solution declined to 57.9±2.6% of the initial amplitude.

The chloride and proton transport pathways in CLC Cl^-^/H^+^ exchangers are partially overlapping at their extracellular part containing the gating glutamate Glu_ext_ [4,30,32]. However, these pathways diverge toward the intracellular protein side [33]. Based on the distinct effects on voltage dependent gating and silent transport cycle probability (see Figure 2), we tested whether the adenine nucleotide regulation is specifically coupled to the separate lower part of the proton transport pathway. For these experiments, we focused on mutant E268Q ClC-5 in which the intracellular proton access is blocked [24,25]. The non-transporting phenotype of the mutant prevents ion current measurements [24,25]; however, gating charge analysis revealed that the nucleotide effects on ClC-5 voltage dependence are abolished. (Supplementary Figure S6). Hence, the adenine nucleotide regulation of voltage-dependent gating depends on the internal (cytosolic) delivery of protons into the CLC transporter machinery.

### Nucleotide chemistry and cytosolic Mg^2+^ are important for the adenine nucleotide regulation of CLC transport

We next performed experiments to investigate the specificity of CLC adenine nucleotide regulation. It was previously reported that the non-hydrolysable ATP analog adenylyl imidodiphosphate (AMP-PNP) cannot augment ClC-4 transport currents [14]. Patch-clamp measurements with 5 mM AMP-PNP in the pipette solution revealed that ClC-5 behaves similarly (Figure 4). The comparable ClC-5 current rundowns observed in the presence of intracellular AMP-PNP and in the presence of AMP (see Fig. 1) suggest that the terminal bridge oxygen of the ATP triphosphate chain is important for the regulation of mammalian CLC transport. In the same set of experiments, we investigated the effects of S-adenosyl-methionine (SAM or AdoMet), a small molecule that activates human cystathionine-beta-synthetases by interacting with their CBS domains [34,35]. In contrast to AMP-PNP, the rundown observed with SAM in the pipette solution resembled nucleotide-free measurements (Figure 4). It seems, therefore, that the binding pocket in the CBS domains of ClC-5 can only accommodate an adenosine moiety containing at least one phosphate group.

**Figure 4.**
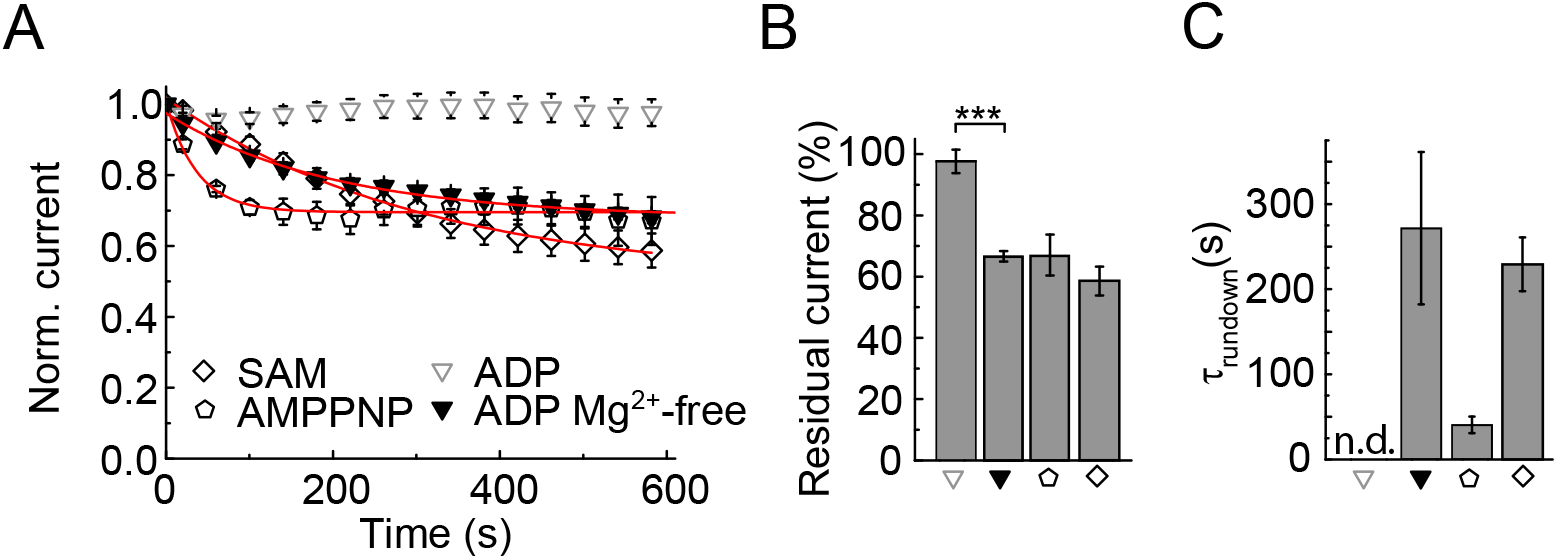
Nucleotide chemistry and Mg^2+^ are important for the regulation of CLC transport. **(A)** Effects of different adenine nucleotides or their washout on the ClC-5 current amplitudes. The data were normalized to the initial current amplitude determined shortly after establishing whole-cell configuration. Some of the data points are omitted for clarity. For comparison, the ADP data set from Figure 1 is also depicted. Red lines indicate monoexponential fits to the data. **(B)** Normalized residual currents determined after 600-s of cell dialysis as shown in (A). **(C)** Averaged rundown time constants obtained from fits with monoexponential decay functions as shown in (A). Asterisks indicate statistically significant differences with p < 0.001 (n=4-7); n.d. – not defined.

In mammalian cells, adenine nucleotides form biologically active complexes with Mg^2+^ [36]. To our knowledge, it has not been investigated whether Mg^2+^ ions play a role in the adenine nucleotide regulation of CLC transport. However, a reduced [Mg^2+^] was used in a previous study [14] in which ADP was found unable to enhance ClC-4 transport. Therefore, we eliminated the MgCl_2_ from our pipette solution. In contrast to experiments with 5mM MgCl_2_ in the pipette (see Figure 1), a significant ClC-5 current rundown was observed after obtaining the whole cell configuration (Figure 4). These experiments demonstrate that Mg^2+^ ions are important for the adenine nucleotide regulation of mammalian CLC transport.

## Discussion

Using mutagenesis, heterologous expression, and electrophysiology, we could demonstrate that cytosolic ATP and ADP enhance the electrogenic ion transport of three broadly expressed mammalian CLC transporter isoforms via binding to the C-terminal CBS domains of these proteins. In contrast, we found that cytosolic AMP inhibits CLC transport (Figures 1 and 2). The role of the CBS structure as energy sensing domain has been demonstrated for a variety of proteins and the AMPK family is probably the most prominent example [12]. By analogy, the differential response to ATP/ADP, on the one hand, and AMP, on the other, equips CLC transporters with a means of sensing and responding to changes in the cellular energy state.

The here-presented findings are in line with previous investigations into the interaction between adenine nucleotides and mammalian CLC transporters [14][20]. They are also coherent with the different effects of AMP and ATP on the *Arabidopsis* AtCLC transporter [37]. Our data are also in agreement with most of the findings of a previous comprehensive investigation on ClC-5; except for the published ion current increase observed in experiments with intracellular AMP (see [15]). It should be noted, however, that the effects analyzed in this particular study [15] were very small and comparable to single channel amplitudes of endogenous ion transporting proteins observed in the same experimental system [38,39]. We suspect, therefore, that such endogenous conductances, probably activated by AMP, might have obscured the CLC current reduction induced by this nucleotide.

Gating current analysis revealed two distinct mechanisms of action by which cytosolic adenine nucleotides regulate mammalian CLC exchangers. On the one hand, we found that adenine nucleotide binding alters the ratio between CLC ion transport and CLC gating charge (Figure 2). Earlier investigations have shown that this ratio depends on the delivery of internal protons into the CLC transporter machinery [17,24,25]. We concluded, therefore, that ATP and ADP binding to the C-terminus of mammalian CLC exchangers facilitates the entry of internal protons into the proton transport pathway whereas AMP binding impedes this entry. This conclusion is supported by the lack of ATP effects in experiments with lower intracellular pH [15] and by the abolished internal pH dependence of mutant ClC-5 E268Q in which the internal proton pathway has been blocked [24,25]. In addition to this effect, we found that cytosolic adenine nucleotides alter the voltage-dependent activation of the investigated CLC isoforms (Figure 2), a process that is governed by the gating glutamate Glu_ext_ [19,24,25,40–42]. The shift resembles the effect observed at lower the cytosolic pH [24,25] and can be similarly explained by an increased delivery of internal protons into the CLC protein. In support of this conclusion, mutation E268Q that blocks the internal entry of the proton transport pathway [17,24,25,28,33] also greatly reduced the effects of adenine nucleotide on the ClC-5 voltage dependence (Suppl. Figure S6).

Especially the results on ClC-3 (see Figures 1 and 2) underline the importance of voltage dependent gating for the ion transport decrease induced by cytosolic AMP. However, as the depolarization-activated gating is right-shifted also by intracellular ATP and ADP (Figure 2), the altered voltage dependence cannot explain the ion transport increase induced by these nucleotides (Figure 1). Therefore, the effects of cytosolic ATP and ADP are rather explained by reduced silent transport cycle probability (Figure 2). This conclusion is in full agreement with the preserved ability of ATP to enhance the ion transport of the Glu_ext_ mutant E211G ClC-5 in which voltage-dependent gating is abolished [31] (see Figure 3). A negatively charged and protonatable residue, such as the native Glu_ext_, is absolutely required for CLC proton transport [26,27,29–31]. Therefore, it might appear surprising at first glance that adenine nucleotide action is preserved in mutant E211G ClC-5. However, it has been established that the Cl^-^ and H^+^ transport pathways of CLC exchangers overlap only partially in their extracellular part that contains the Glu_ext_ but diverge toward the intracellular protein side [33]. Consequently, neutralizing the Glu_ext_ should not affect the entry of internal protons into the separate lower part of the transport machinery. Along the same line of reasoning, CLC channels, while unable to transport protons, have preserved parts of the proton transport machinery of the archetypic CLC Cl^-^/H^+^ exchangers [43]. The here investigated ClC-5 E211G abolishes proton transport and converts ClC-5 into a passive Cl^-^ conductor resembling the classical CLC channels [31]. Therefore, the strong cytosolic pH dependence of the ATP regulation of the ClC-1 channel [44,45], together with the preserved ATP action on the ClC-5 transporter Glu_ext_ mutant (E211G ClC-5, see Figure 3), both hint at the existence of common molecular mechanisms underlying the adenine nucleotide regulation of CLC channels and transporters.

What are the molecular mechanisms that link the CBS domain nucleotide binding to CLC proton transport? Whereas we cannot directly compare the ATP-bound and the ATP-free conformations of the CLC C-terminus [6], it is established that the intracellular entrance of the CLC proton transport pathway is localized in the proximity of the region connecting the C-terminus and the CLC transmembrane core [25,33,46,47]. It is also established that ATP binding to the ClC-5 CBS domains induces large-scale conformational changes of the whole CLC C-terminus [13]. Such rearrangements are likely to be propagated towards the transmembrane core and to affect the injection of protons into the CLC transporting machinery. This notion is supported by the significant impact that both the C-terminus and internal protons exert on the voltage dependent gating of various mammalian CLC transporters [19,24,25,49]. Additional support is provided by the recently demonstrated long-range communication between the C-terminus and the CLC transmembrane core of the anion channel homolog CLH-3b [48].

Our experiments revealed that adenine nucleotide action depends on the cytosolic availability of Mg^2+^. It has been previously shown that the ClC-5 CBS domains can bind ADP also in the absence of Mg^2+^ [6]. As such binding does not increase CLC ion transport (Figure 2), it is likely that Mg^2+^ ions fine-tune the conformational changes that lead to full transport activation. At this point, we can only speculate about the physiological role of this fine-tuning mechanism, and in general about the role of the CLC transporter adenine nucleotide regulation. However, it is striking that both the voltage dependence and the silent transport cycle probability are affected by pathogenic ClC-5 mutations associated with Dent disease [46]. As vesicular secondary active exchangers [17,26,27], ClC-3, ClC-4 and ClC-5 utilize the electrochemical gradient built up by ATP-dependent pumps [2]. Hence, an [AMP] increase upon depletion of the cellular chemical energy sources will suppress coupled CLC transport and reduce ATP consumption. In addition, the associated higher silent transport cycle probability will increase the vesicular membrane capacitance [17,25] and reduce the energy barrier opposing the V-ATPase-mediated vesicular acidification [18]. Drawing an analogy to the proteins of AMPK family [12], our findings suggest that mammalian CLC exchangers might participate in a survival program activated during cell starvation or other extreme conditions. Based on the expression pattern and vesicular localization of the here-investigated isoforms (recently reviewed in [2]), the metabolic energy dependence of intracellular CLC transport might become highly relevant during various ischemic, hypoxic, or anoxic conditions of the kidney, heart, and brain, including acute injuries, diabetes, heart attack, stroke, etc. [50–56]. The selective apoptosis of hippocampal neurons in ClC-3 knockout mice [57] might reflect a higher susceptibility to cell death induced by energy deprivation. ClC-3 sensing of the ATP/ADP/AMP ratio might control insulin production by defining the rate of granular acidification [58]. Pathogenic ClC-4 loss-of-function mutants [59,60] might reduce the ability of neurons to react to metabolic stress, and accordingly might affect cognitive function.

Whereas the physiology of adenine nucleotide regulation remains to be investigated, the intracellular localization of ClC-3, ClC-4 and ClC-5, together with their pivotal role in endosomal and lysosomal function, renders such regulation physiologically meaningful and creates a link between the cellular metabolic state and the activity of the endolysosomal system.

## Materials and Methods

### Cell culture, construct generation, expression

DNA sequences encoding for the investigated isoforms ClC-3, ClC-4 and ClC-5 were inserted into specially functionalized expression vectors and controlled by sequencing. For ClC-4, the bicistronic pRcCMV vector contained additionally the CD8 surface antigen, translationally controlled by an internal ribosome entry site (IRES [61]) to allow identification of successfully transfected cells with anti-CD8 antibody-coated polystyrene beads (Dynabeads CD8; Gibco, Life Technologies, Darmstadt, Germany). For the expression of WT ClC-5, the DNA sequences of the RFP variant mCherry and the CLC protein were fused together into the pRcCMV expression vector (described elsewhere [25]), allowing the selection of transfected cells by fluorescence. The used ClC-3c eGFP or mRFP constructs into the FsY1.1 G.W. or p156rrL vectors are described elsewhere [17]. Point mutations for amino acid substitutions or truncations were generated using the QuikChange site-directed mutagenesis kit (Stratagene, La Jolla, CA, USA).

HEK293T cells were transiently transfected with 5 – 10 μg plasmid DNA using calcium phosphate precipitation [62]. Cells were cultured in DMEM (Gibco, Life Technologies, Darmstadt, Germany), supplemented with 10% fetal bovine serum (Biochrom, Berlin, Germany), 2 mM L-glutamine and 50 unit/ml penicillin/streptomycin (Sigma-Aldrich Chemie, Munich, Germany). In some cases, previously described stable HEK293 cell lines expressing WT ClC4 or WT ClC-5 [19,25] were used for electrophysiology; stable expression was maintained using 0.9 mg/ml Geneticin (G418) added to the standard HEK293 growth medium (MEM, supplemented with 10% FBS, both from Gibco, Life Technologies, Darmstadt, Germany).

### Electrophysiology

Whole-cell or inside-out patch-clamp measurements [63] were performed 48 h posttransfection using a HEKA EPC-10 amplifier and PATCHMASTER software (HEKA Electronics, Lambrecht, Germany), or an Axopatch 200B amplifier with Clampex software (Molecular Devices, Sunnyvale, CA, USA). Filtering with 3 to 10 kHz was applied prior to digitizing the data at a 30-kHz sampling rate. Patch pipettes from borosilicate glass were fabricated with a P-97 puller (Sutter Instrument, Novato, CA, USA) and heat-polished to reach a resistance between 0.9 and 2 MΩ on an MF-900 Microforge (Narishige, London, UK). If required (i.e. for inside-out measurements), pipettes were coated with dental wax to minimize their electric capacitance. Series resistance compensation and capacitive cancellation minimized the voltage error to <5mV. Linear capacitive artifacts were further reduced by a P/4 leak subtraction sequence [64] applied from a holding potential of −30 mV. The standard extracellular solution contained (in mM) 145 NaCl, 15 HEPES, 5 MgCl_2_, 4 KCl, 1 CaCl_2_. The nucleotide-free pipette solution contained: (in mM) 120 NaCl, 15 HEPES, 5 MgCl_2_, 1 EGTA. Adenine nucleotides (5 mM Na-ATP, 5 mM Na-ADP or 5 mM Na-AMP, all from Sigma-Aldrich) were added to the pipette solution shortly before the experiments and the pH was additionally adjusted if necessary. All solutions were titrated to pH 7.4 using NaOH. In NMDG-based solutions, NaCl was substituted by equimolar N-methyl-d-glucamine-Cl; pH was adjusted with NMDG.

### Gating current measurements to estimate the apparent CLC gating charge

The gating charge mobilized during CLC activation was obtained by integrating the area under P/4 subtracted off-gating currents elicited by voltage jumps in the dynamic range of the investigated CLC transporter [24]. For further analysis, the gating charges were normalized either to the maximal gating charge at saturating voltages or to the steady-state the ionic current at +165 mV.

To describe the voltage dependence of the charge movements (*Q(V)*), the following standard Boltzmann function (Equation 1) was fitted to the data:

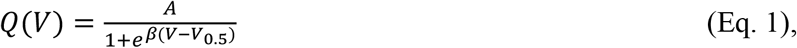

with:

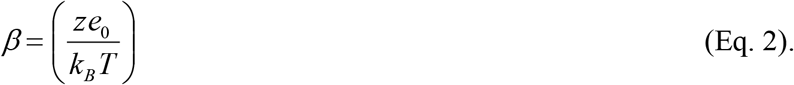

Here, *V*_*0.5*_ and *z* denote the voltage for half-maximal activation, and the apparent number of elementary charges *e*_*0*_ displaced in the transmembrane electric field during the investigated voltage-dependent transition. The Boltzmann constant and the absolute temperature are indicated as *k*_*B*_ and *T*, respectively. The apparent amplitude of the Boltzmann function is indicated as *A*.

### Structural modeling and Data analysis

The UCSF Chimera [65] interface to MODELLER [66] was used to create a 3D protein homology model of the full-length ClC-5 amino acid sequence based on the crystallized structures of the soluble CBS-domain of ClC-5 (PDB-ID: 2J9L [6]) and the transmembrane domains of CmCLC (PDB ID: 3ORG [5]). Experimental data were analyzed using a combination of FitMaster (HEKA) or Clampfit (Molecular devices), Excel (Microsoft) and Origin (OriginLab Corporation, Northampton MA, USA). Statistical significance was assessed using a two-sample t-test. All summary data are shown as mean ± SEM.

## Acknowledgments

We thank Petra Killian, Birgit Begemann and Toni Becher for technical assistance.

## Author contributions

MG, KL, RG, ChF, and AKA contributed to the design of the work. MG, KL, RG, and AKA, and AKA contributed to acquisition and analysis of the data. AKA drafted the manuscript. MG, KL, RG, ChF, and AKA revised the paper critically for important intellectual content.

